# Multi-scale Assessment of Brain Blood Volume and Perfusion in the APP/PS1 Mouse Model of Amyloidosis

**DOI:** 10.1101/2022.07.01.498298

**Authors:** Leon P Munting, Marc PP Derieppe, Lenard M Voortman, Artem Khmelinskii, Ernst Suidgeest, Lydiane Hirschler, Emmanuel L Barbier, Baudouin Denis de Senneville, Louise van der Weerd

**Affiliations:** Leiden University Medical Center, Leiden, the Netherlands; Prinses Máxima Center for Pediatric Oncology, University Medical Center Utrecht, Utrecht, the Netherlands; Netherlands Cancer Institute, Amsterdam, the Netherlands; Université Grenoble Alpes, Grenoble, France; Université de Bordeaux, Bordeaux-Sud-Ouest, France

## Abstract

Vascular dysfunction is increasingly recognized to play a role in the development of Alzheimer’s disease (AD). The relation between vascular dysfunction and the neuropathological amyloid β accumulation characteristic for AD is however unclear. The limited resolution of in vivo imaging techniques, the intricate 3D structure of the microvasculature and the different co-occurring types of amyloid β accumulation in patients hamper studying this relation in patients. Here, we therefore employed the APP/PS1 mouse model, which develops parenchymal amyloid β plaques, to study the effect of parenchymal amyloid β plaques on the structure and function of the vasculature. Blood vessels and amyloid β plaques were fluorescently labeled in vivo with lectin-DyLight594 and methoxy XO4, respectively, in APP/PS1 mice at old age. The brain tissue was cleared post-mortem with the CUBIC clearing protocol, which allowed structural imaging at microscopic resolution of the vessels and plaques in a large 3D volume. Segmentation of the vasculature enabled mapping of the microvascular Cerebral Blood Volume (mCBV), which ranged from 2 % to 5 % in the white matter and the thalamus, respectively. No mCBV differences were observed between APP/PS1 mice and wild type (WT) control mice. The effect of the amyloid β plaques on vascular function was studied in vivo by measuring Cerebral Blood Flow (CBF) and Arterial Transit Time (ATT) with Arterial Spin Labeling (ASL) MRI. Similar to the mCBV findings, no differences were observed in CBF or ATT between APP/PS1 and control mice, indicating that brain vascular morphology and function in this mouse model are preserved in the presence of amyloid β plaques.

## INTRODUCTION

Familial mutations in genes related to the production of the amyloid-β peptide are associated with early cognitive decline and extensive accumulation of amyloid-β pathology in the brain. These mutations lead to Alzheimer’s disease (AD) with full penetrance [1]. This is a clear indication that amyloid-β plays an important role in the development of AD, at least in familial cases with early onset. The importance of its role in the development of sporadic AD, however, is continuously debated. Researchers who are sceptic about the amyloid hypothesis, point out that amyloid-β plaque load correlates poorly with cognitive decline and that plaques are also found in the healthy elderly [2]. Alternative, or synergistic, pathogenic mechanisms for AD are receiving more and more attention in the recent years, among which is the cerebral circulation. Various imaging studies have demonstrated that changes in brain perfusion [3], blood volume [4,5] and cerebrovascular reactivity [6] are associated with AD. In a predictive model based on human imaging data, cerebrovascular dysfunction has been estimated to be present early on in the AD onset [7]. AD and cardiovascular disease share genetic risk factors [8] and histological studies have shown that cerebrovascular pathology such as microinfarcts and atherosclerotic arteries is commonly found in AD, more often than in other neurodegenerative diseases [9]. Given the lack of evidence to hold either amyloid-β or the vasculature fully responsible for the development of sporadic AD, it seems likely that it is a multifactorial disease in which amyloid-β, cerebrovascular dysfunction and possibly other factors all play a part. Insight into the extent of their roles as well as how amyloid-β and cerebrovascular dysfunction influence each other is important to better understand the mechanisms of the disease and for the development of therapeutic strategies. However, studying this relation in patients is complicated due to a number of reasons. Firstly, although PET and MRI imaging techniques exist to directly visualize amyloid-β burden and perfusion *in vivo* in patients [1,3], these imaging techniques are not offering sufficient spatial resolution to directly link amyloid-β distribution and the microvasculature. Thus, these cellular relations can only be characterized end-stage after autopsy. Secondly, the intricate 3D structure of the microvasculature complicates its characterization even *ex vivo*, as conventional histology does not allow volumetric imaging. Lastly, amyloid-β can be present in different forms, including parenchymal amyloid-β plaques, cerebrovascular amyloid-β and amyloid-β oligomers. The latter are soluble, neurotoxic forms of amyloid-β aggregates which in the recent years have been shown to correlate better with cognitive decline than parenchymal amyloid-β plaques [10]. Cerebrovascular amyloid-β is commonly referred to as cerebral amyloid angiopathy (CAA), which has been estimated to co-occur in around 78 % to 98 % of the AD cases [11]. A deeper understanding of the pathophysiology implies that the interplay of each structure, *i*.*e*., amyloid-β oligomers, cerebrovascular and parenchymal amyloid-β, with the brain vasculature is better understood.

Mouse models can be a useful tool to study the relation between cerebrovascular dysfunction and amyloid-β, as tissue can be collected for histology at any age and disease stage following *in vivo* imaging in order to relate imaging outcomes with molecular or microstructural phenotypes. Furthermore, the possibility to specifically engineer the genome of a mouse has enabled the creation of models that reflect separate disease components. By introducing different types of familial AD mutations, models have been created that develop either mainly parenchymal plaques [12,13], arteriolar CAA [14], capillary CAA [15] or oligomers [16]. Here, we exploited this benefit of mouse models and studied whether the presence of parenchymal amyloid-β plaques influences the morphology or function of the brain vasculature. Specifically, we studied the microvascular cerebral blood volume (mCBV), the cerebral blood flow (CBF) and the arterial transit time (ATT) in the APP ^swe^/PS1^dE9^ amyloidosis model [12] at old age and thus with high parenchymal plaque burden. To measure the mCBV, the vasculature was imaged post-mortem in 3D with laser scanning microscopy after the brain tissue was made transparent with CUBIC (clear, unobstructed brain imaging cocktails and computational analysis)[17]. CBF and ATT were measured *in vivo* with pseudo-Continuous Arterial Spin Labeling (pCASL)-MRI. Our findings indicate that the structure and function of the mouse cerebral circulation are not substantially disturbed by the presence of parenchymal amyloid-β plaques.

## MATERIALS & METHODS

### Animals

This study was carried out in compliance with the European Directive 2010/63/EU on the protection of animals used for scientific purposes and reported in compliance with the ARRIVE guidelines [18]. The experiments were approved by the institution for animal welfare of the Leiden University Medical Center in the Netherlands and performed under DEC12065. The background strain of all mice in the study was the F2 generation of a cross between C57Bl/6J and C3H/HeJ mice. An optical imaging study and an MRI study were performed with two separate groups of mice bred in-house. The founder mice were acquired at the Jackson Laboratory (Bar Harbor, ME, USA). For optical imaging, 5 APP s^we^/PS1^dE9^ (APP/PS1)[12] transgenic (TG) mice (17 ± 2 months old; 2 female) and 5 wild type (WT) controls (18 ± 2 months old; 3 female) were used. In the MRI study, 6 APP/PS1 mice (24 ± 1 month old; 6 females) and 6 WT controls (26 ± 1 month old; 4 females) were used. The mice were housed together (2–4 per cage) and had access to chow food and water *ad libitum*. The cages were individually ventilated and were provided with bedding material and cage enrichment, with a 12 h day/night cycle.

### Microscopy

#### Brain isolation and tissue processing

Twenty-four hours ante-mortem, the 5 APP/PS1 and 5 WT mice were i.p. injected with 10 mg/kg methoxy-XO4 (in-house synthesized, 0.05 M dissolved in 1:1 DMSO:Cremophor and diluted in PBS to a total volume of 150 μL), a small fluorescent molecule that binds to beta-sheets and thereby enables visualization of Aβ deposits. The next day, 5 minutes ante-mortem, the mice were i.v. injected with 200 μL of 1 mg/mL DyLight 594-labeled tomato lectin (Vector Labs, CA, USA), which binds to the endothelium and thereby enables visualization of the vasculature. Subsequently, the mice were injected with an i.p. overdose of Euthasol (AST Pharma), followed by opening of the chest for intra-cardiac perfusion with 20 mL of ice-cold PBS and 20 mL of ice-cold 4 % PFA. The brains were then isolated and fixated further in 4 % PFA. Thereafter, the brain was cut in two hemispheres. The left hemisphere was made transparent for imaging, the right hemisphere was stored overnight in 30 % sucrose in PBS, snap-frozen in 2-methylbutane and stored at -80 °C for later studies. To make it transparent, the left hemisphere was treated with the CUBIC clearing protocol, which is described in [19]. In short, the hemisphere was washed 2 times for 1 hour in PBS at room temperature (RT) to wash away the PFA, transferred to a 1:1 mix of PBS and CUBIC solution 1 (25 % urea, 25 % quadrol [N,N,N⍰,N⍰-Tetrakis(2-hydroxypropyl)ethylenediamine, Sigma-Adrich] and 15 % triton weight/weight in MiliQ) and incubated with gentle shaking for 6 hours at 37 °C. Thereafter, the 1:1 mix was replaced with pure CUBIC solution 1 and incubated for 8 days at 37 °C with gentle shaking. During this time, the solution was replaced every 2 days. Afterwards, the CUBIC solution 1 was washed away with 6 PBS washes over the course of 6 hours and the brain tissue was then incubated overnight at RT with a 1:1 mix of PBS and CUBIC solution 2 (50 % sucrose, 25 % urea and 10 % TEA [Triethylamine, Sigma-Aldrich] weight/weight in MiliQ). The next morning, the mix of PBS and solution 2 was replaced with pure CUBIC solution 2, and further incubated for 2 days with gentle shaking at 37 °C. The solution was refreshed once after 1 day. The result of the clearing procedure in terms of tissue transparency is illustrated in figure 1.

**Figure 1:**
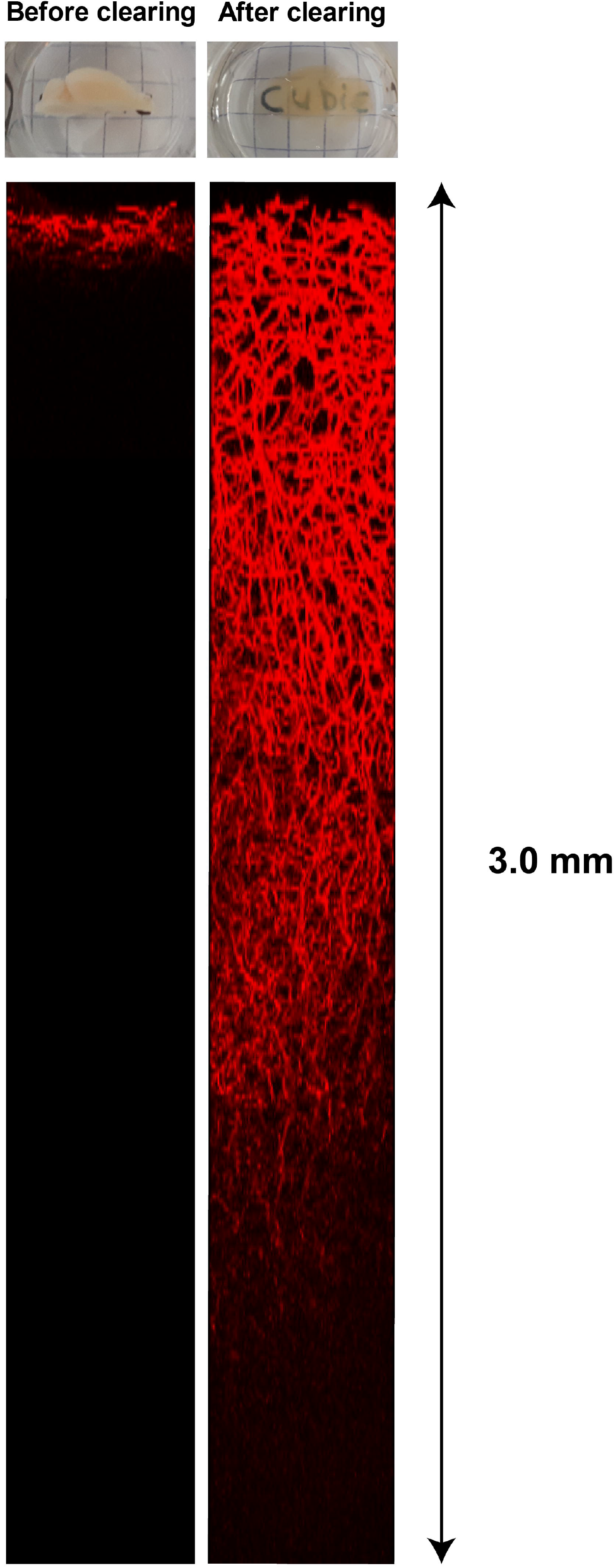
Effect of CUBIC clearing on tissue transparency and penetration depth with imaging. On the top left, a hemisphere is shown before clearing; on the top right the same hemisphere after clearing. On the bottom left, an orthogonal view from a representative Z-stack is shown, acquired in an uncleared hemisphere. On the bottom right, an orthogonal view from a representative Z-stack acquired in a cleared hemisphere, with similar acquisition parameters. Note the increase in penetration depth when imaging cleared tissue. The Z-stacks were acquired at the level of the thalamus, in sagittal orientation, without laser power compensation over imaging depth.

#### Image acquisition

The hemisphere was imaged in sagittal orientation in a 1:1 mix of silicone oil (Sigma, product number 175633) and mineral oil (Sigma, product number M8410) with a Zeiss LSM 710 NLO laser scanning microscope. A Zeiss 20x Clr Plan-Apochromat objective (NA = 1.0; WD = 5.6 mm; Corr nd = 1,38) and a motorized stage were used, allowing imaging deep into the tissue together with full-brain coverage in the XY-direction. The methoxy-XO4 and DyLight594 were simultaneously excited with a multiphoton Ti:Sapphire laser (Mai Tai, Spectra Physics) at 800 nm and their emission was filtered and detected with a Zeiss non-descanned Binary GaAsP PMT detector (BiG.2) equipped with two filters (green 500–550 nm and red 570–610 nm). Even though the hardware of the microscope allowed imaging with sub-micrometer axial resolution, the axial resolution was limited to 3.32 μm, with a 9.96 μm step size. This was done to find a trade-off between data quality on the one hand and acquisition time and file size on the other hand, which were around 2.5 days and 10 GB per sample, respectively.

#### Image processing

The resulting Z-stacks were flat-field corrected using a custom macro in ImageJ and stitched with the Grid/Collection plug-in [20] in ImageJ, using the stage coordinates as initialization. Stage coordinates were extracted from the metadata of the Zeiss (.czi) file using a custom Python script. Methoxy-XO4 has a very wide emission spectrum, therefore amyloid plaques were visible in both the green and red channel (data not shown). DyLight-594 has a narrow emission spectrum and was thus visible in the red channel only. To remove the methoxy-XO4 fluorescence from the red channel, the green channel was first subtracted from the red channel, also for the WT mice for the sake of comparability. Vessel segmentation was performed with the subtracted data using an Otsu threshold. One mouse (male, WT) had to be excluded from further analysis, due to wrong segmentation, which was the result of a low SNR in the raw data, probably a result of inadequate i.v. lectin injection. After segmentation, the blood volume in the brain was mapped using patches of 15 × 15 × 5 voxels in which the volume percentage of vessel-segmented voxels was determined, resulting in mCBV maps with 50 μm^3^ isotropic resolution. To retrieve Atlas-based regions of interest (ROIs), the stitched and shading-corrected data (red channel only, before subtraction) was downsampled to 50 μm^3^ isotropic resolution and registered to the Allen Brain template using EVolution, an edge-based variational non-rigid multi-modal image registration method [21]. Six different ROIs were chosen from the Allen Brain template and propagated to the blood volume maps of each individual mouse. This process was verified and adjusted if needed by a user (LPM). If a large vessel was present in a brain region, this resulted in patches with a high blood volume (> 15 %, see red arrow in figure 2). To avoid such macro-vascular contribution to the regional blood volume values, only voxels with values below 15 % of blood volume were included in the analysis. The impossibility to place the brain tissue exactly orthogonal below the microscope objective resulted in oblique optical sectioning (supplementary figure 1). Also, decreasing signal intensity values were observed when deeper parts of tissue were imaged, even though the brain tissue was CUBIC cleared (figure 1). This resulted in decreasing mCBV values over imaging depth. Together with the oblique optical sectioning, this led to mCBV maps with an axial gradient due to differing quantities of tissue present above different tissue regions in one slice (supplementary figure 1). Alignment of the tissue in the Z-direction with a custom MATLAB script removed this axial gradient, but maintained the gradient in the Z-direction, starting after around 10 slices/500 μm (supplementary figure 2). The analysis was therefore restricted to the upper 500 μm of tissue.

**Figure 2:**
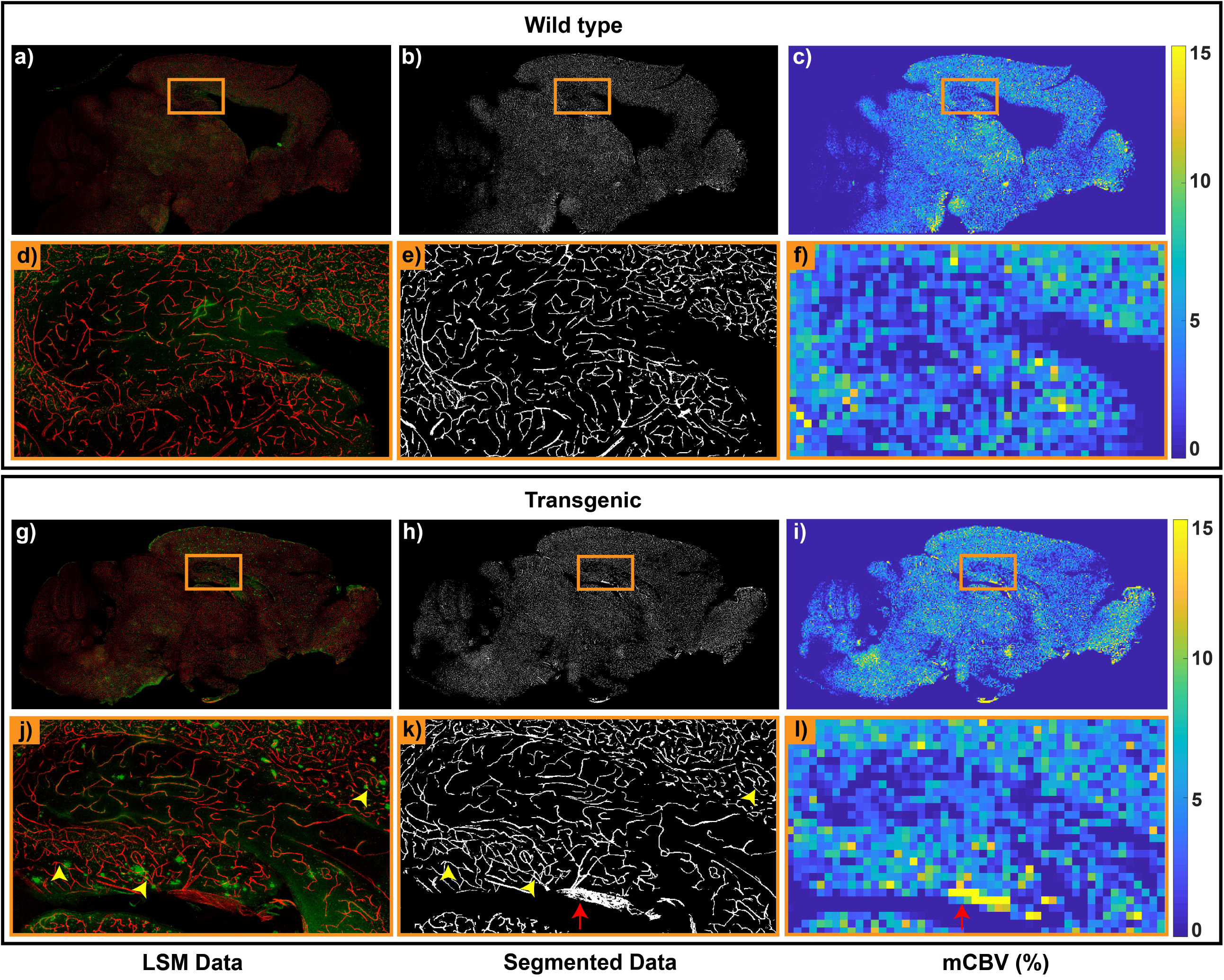
Example of the segmentation and microvascular Cerebral Blood Volume (mCBV) calculation process. A wild type (WT) example is depicted in the upper two rows (a-f), an APP/PS1 transgenic example in the lower two rows (g-l). The images on the second (d-f) and fourth (j-l) rows correspond to the orange squares in the images on the first and third rows. In the images in the left column (a,d,g,j), maximum intensity projections of 5 raw data slices are shown. In the middle column, the result of the Otsu segmentation and on the right column, the resulting mCBV maps. The yellow arrowheads in (j) and (k) point to vessels that are in the close vicinity to plaques, but seem unaffected by them. The red arrow in (k) and (l) points to a large vessel, which results in very high values in mCBV map.

### MRI

The 6 APP/PS1 and 6 WT control mice were scanned with a Bruker 7 T PharmaScan (Ettlingen, Germany) and a 23 mm volume coil, using the same protocol as described in [22]. The 6 WT mice were the same as used in [22]. In short, the following protocol was used:

#### Animal preparation

the mice were anesthetized with 3.5 % isoflurane (Pharmachemie BV, Haarlem, Netherlands) for induction and 1.5-2.0 % for maintenance in a 1:1 mix of air and oxygen, to keep breathing rates around 100 bpm. The temperature was kept at 37 °C with a feedback-controlled waterbed (Medres, Cologne, Germany).

#### Image acquisition

Anatomical T_2_-weighted (T2W) images were first acquired in all three orientations with a standard Bruker T_2_RARE sequence for planning the ASL sequences and for registration purposes. Thereafter, the pCASL interpulse phase-increase was optimized with pre-scans, as described before [23]. Then, two pCASL sequences were acquired, one optimized for measuring CBF (standard pCASL), the other for measuring ATT (time-encoded pCASL [te-pCASL]). The standard pCASL consisted of 60 pairs of label and control scans with a labeling duration of 3000 ms, a constant post-labeling delay (PLD) of 300 ms and a total scan time of 7 minutes. The te-pCASL consisted of a 45 times averaged Hadamard12-encoded labeling block with a sub-bolus labeling duration of 50 ms and 11 effective PLDs (30 ms till 530 ms with 50 ms spacing per PLD) and a total scan time of 7 minutes. Both pCASL scans were followed by a 3-slice coronal spin echo EPI readout with identical parameters (17 ms echo time, 0.224 × 0.224 mm^2^ in-plane resolution, 1.5 mm slice thickness and a 1 mm slice gap). To aid quantification, an inversion-recovery scan and a flow-compensated, pCASL-encoded FLASH sequence were additionally acquired for every mouse to measure the tissue T_1_ and the labeling efficiency, respectively. One mouse (TG, female) had to be excluded from further analysis, as it died during the scan, probably a result of fragility due to old age.

#### Image processing

The individual EPIs from both ASL sequences were first aligned to the first EPI of the standard pCASL scan to compensate for motion during the scans. Six different ROIs (auditory/visual cortex, sensory cortex, motor cortex, hippocampus, thalamus and striatum) were manually drawn on the anatomical images of one arbitrarily chosen mouse (reference dataset), and propagated to the anatomical images of the other animals after registration to the reference dataset. The anatomical images, T_1_ maps and te-pCASL scans of all mice were registered to the first EPI of their standard pCASL scan, allowing the propagation of the ROIs to all MRI sequences. All steps were performed in a coarse-to-fine manner using the Elastix image registration toolbox [24] and verified by two independent observers. Buxton’s kinetic perfusion model [25] was used to calculate the CBF and ATT. The CBF was derived from the averaged signal difference between the label and control (ΔM) scans from the standard pCASL sequence. The ATT was retrieved after fitting the 11 effective PLDs from the te-pCASL sequence to the perfusion model.

### Statistics

For both the microscopy and the MRI data, a mixed ANOVA was used to test whether there was a significant effect of either the genotype or the brain region on the outcome measures mCBV, CBF and ATT. Because Mauchly’s test of sphericity was significant in the CBF data, the degrees of freedom for the F-distribution were Greenhouse-Geisser corrected. For ATT and mCBV data, Mauchly’s test was not significant, thus sphericity was assumed for the degrees of freedom of the F-distribution. Statistical testing was done using IBM SPSS statistics 23 software (Armonk, New York, NY, USA).

## RESULTS

### mCBV mapping

A clear improvement in tissue transparency and imaging depth was seen after brain tissue was treated with CUBIC (figure 1). Representative WT and TG examples of cerebrovascular and amyloid-β plaque labeling are shown in the left column in figure 2, confirming that plaques are present in TG mice only. In the vicinity of these plaques, no noticeable morphological changes could be observed in the capillary bed at the spatial resolution set, see vessels indicated with arrowheads in 2d and additional examples in supplementary figure 3. For the quantitative analysis of the mCBV in the volume of hemisphere imaged, the cerebrovascular signal was first segmented with the application of an Otsu threshold (middle column in figure 2). Thereafter, the density of positively segmented voxels was mapped in patches of 50 μm^3^ isotropic resolution (right column in figure 2). This enabled comparison of blood volumes between WT and TG mice. In figure 3, the first slice of the aligned mCBV maps of all the mice in the study are shown. mCBV estimates were retrieved from 6 different brain regions after registration to the Allen Brain atlas and ranged between 2-5 % (figure 4). No significant difference in mCBV was found between the APP/PS1 and the WT (F[1,7] = 166; p = 0.40), but there was a significant effect of brain region on mCBV (F[5,35] = 43.2; p < 0.001), with the highest blood volume in the thalamus (4.9 % average in WT), and the lowest in the white matter (2.0 % average in WT).

**Figure 3:**
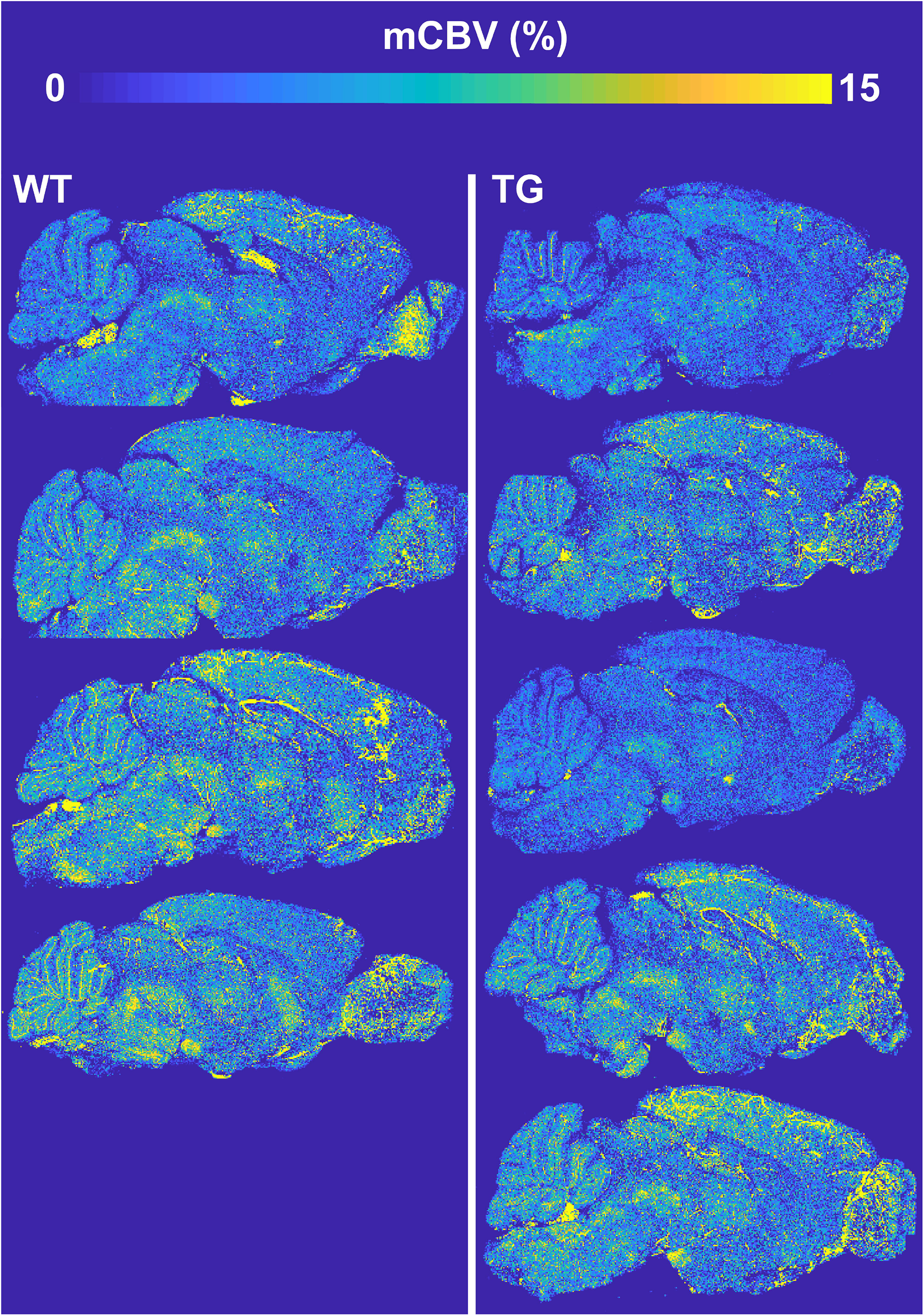
Microvascular Cerebral Blood Volume (mCBV) maps of all mice in the study. The left column represents wild type (WT) mice, the right column transgenic (TG). All maps represent the most superficial mCBV map with respect to the microscope objective.

**Figure 4:**
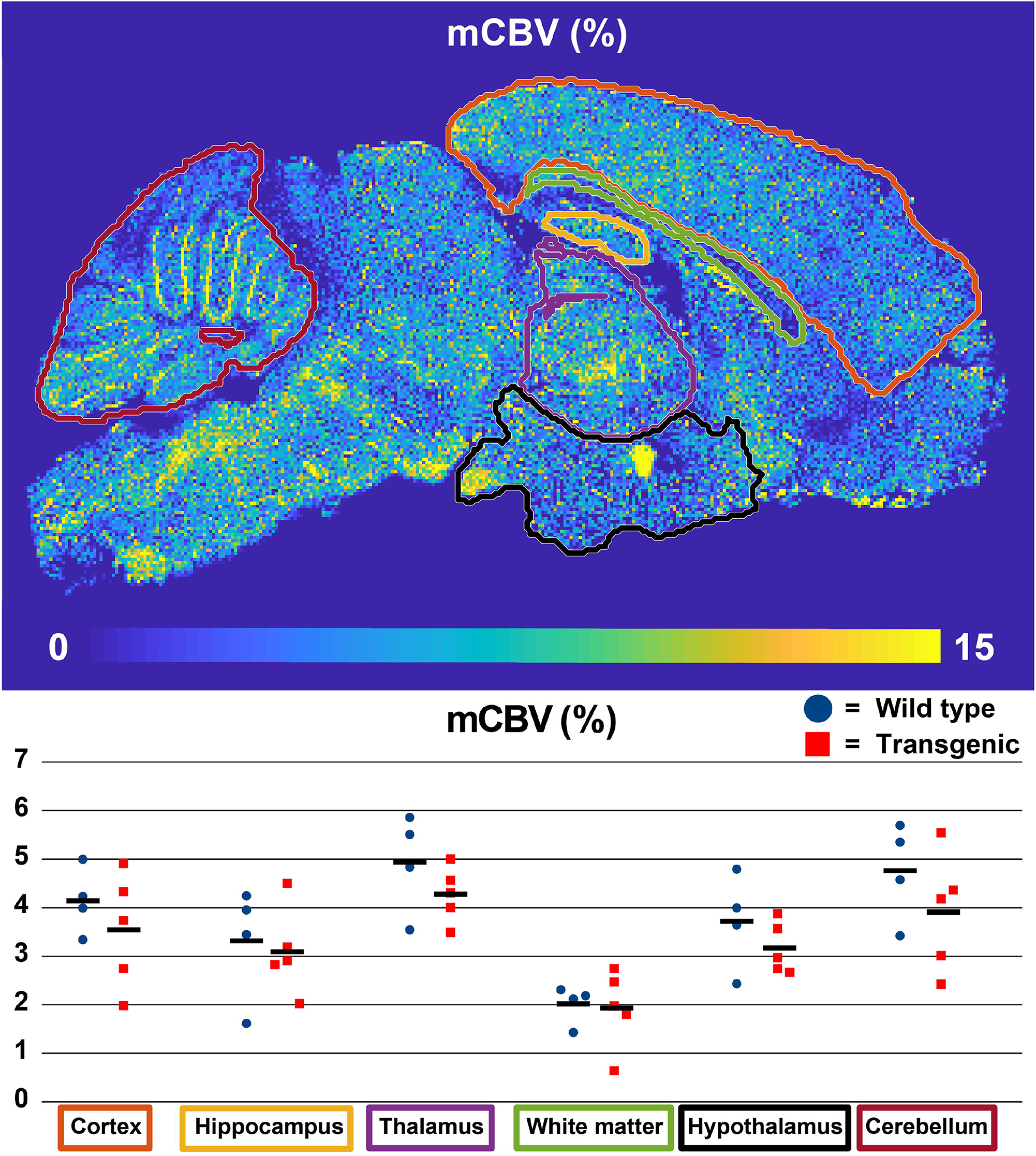
Microvascular Cerebral Blood Volume (mCBV) quantification. mCBV is quantified in 6 brain regions and full brain. Circles (WT) and squares (TG) represent individual mice, stripes represent group averages.

### CBF and ATT mapping

Characteristic CBF maps for both WT and TG mice are shown in figure 5a. The CBF was compared between WT and TG mice in the 6 different brain regions as displayed on an anatomical T2W image in figure 5b. Representative ATT maps for both genotypes are shown in figure 6a. ATT was also compared between WT and TG mice in the same brain regions. No differences were observed in either CBF or ATT between the APP/PS1 mice and the WT mice (F[1,9] = 2.06; p = 0.19 & F[1,9] = 0.056; p = 0.82 respectively). Similar as in the mCBV data, there was a significant effect of brain region on both CBF and ATT (F[1.70,15.0] = 4.62; p = 0.031 & F[5,45] = 84.2; p < 0.001 respectively).

**Figure 5:**
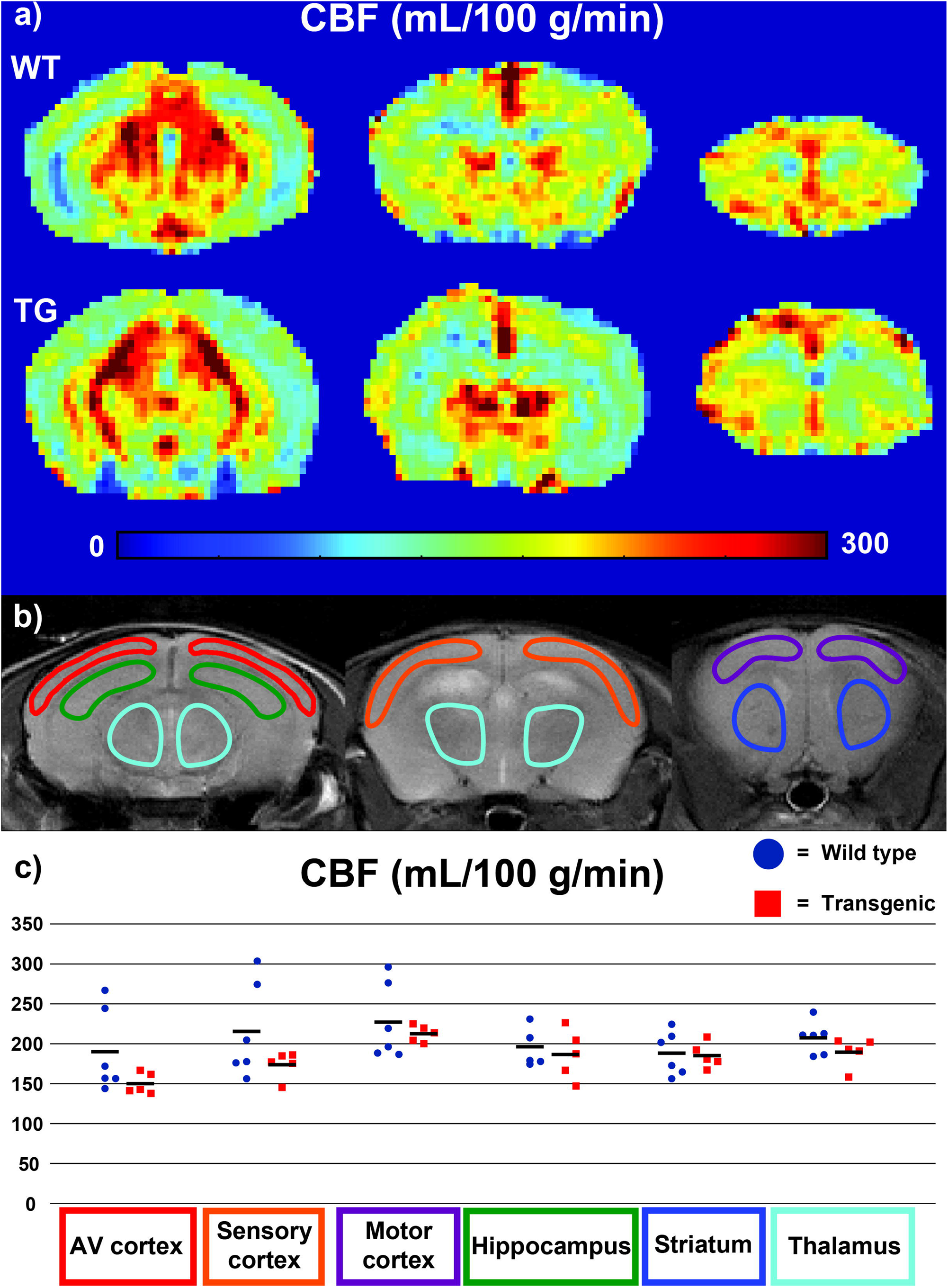
Cerebral blood flow (CBF) maps and CBF values from the MRI data. (a)—CBF maps (three slices) are shown from an average wild type (WT) mouse on the upper row and an average transgenic (TG) mouse on the second row. (b)—Spatially corresponding anatomical images are shown, with the ROIs indicated in which CBF was quantified. (c)—The resulting CBF values are shown in a graph, where circles (WT) and squares (TG) represent individual mice and stripes represent group averages. AV = auditory/visual.

**Figure 6:**
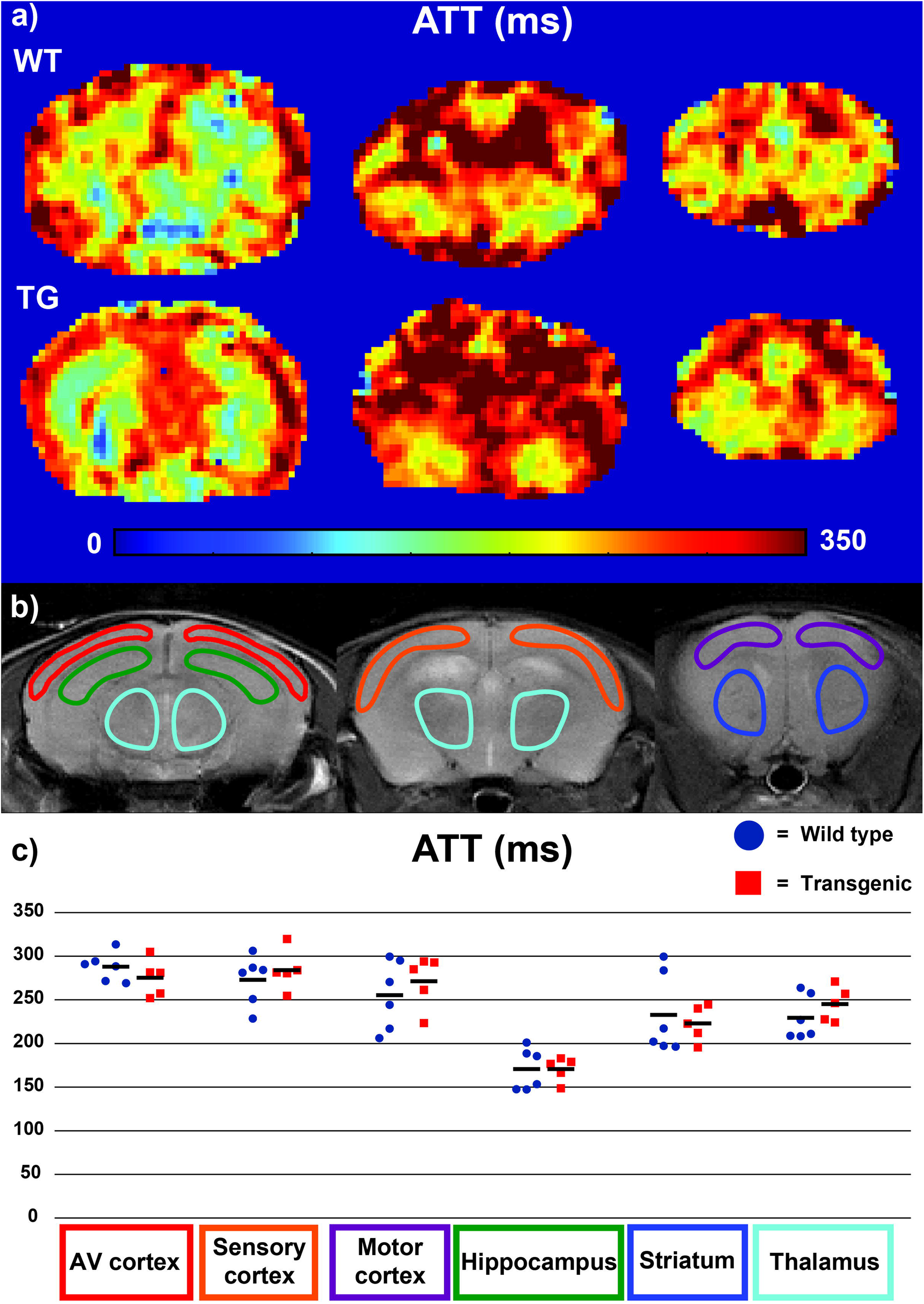
Arterial Transit Time (ATT) maps and ATT values from the MRI data. (a)—ATT maps (three slices) are shown from an average wild type (WT) mouse on the upper row and an average transgenic (TG) mouse on the second row. (b)—Spatially corresponding anatomical images are shown, with the ROIs in which ATT was quantified. (c)—The resulting ATT values are shown in a graph, where circles (WT) and squares (TG) represent individual mice and stripes represent group averages.

## DISCUSSION

Alzheimer’s disease is characterized by the presence of neuropathological amyloid-β accumulations and is associated with cerebrovascular dysfunction [2,7]. The exact contributions of both on AD onset and their interplay are currently not well known and are difficult to study in humans. Mouse models of disease can provide valuable information, as microstructural tissue characterization with histology can be performed following *in vivo* imaging. Also, the many different mouse models reflecting different characteristics of AD allow studying the relation between different aspects of the disease. Here, we performed *in vivo* MR imaging and post-mortem microscopy in a large 3D volume in the APP/PS1 amyloidosis model, which develops extensive amyloid-β plaque accumulation in the brain parenchyma and to a lesser extent CAA on the leptomeningeal arteries [26]. The CUBIC clearing protocol together with the implemented image processing pipeline allowed characterizing amyloid-β plaques and the microvasculature in 3D, including the quantification of the microvascular blood volume in a brain slab of 0.5 mm and direct observation of the effect of plaques on the vasculature. *In vivo*, the perfusion parameters CBF and ATT were measured using ASL-MRI. Given the many connections that have been made between amyloid-β and cerebrovascular dysfunction, it seems remarkable that in this study, no cerebrovascular dysfunction was observed using two imaging modalities in aged APP/PS1 mice with high amyloid burden.

In contrast to our finding, a reduction in CBV has been reported in AD patients compared to non-demented elderly controls [4,5]. A direct capillary loss might underlie this reduction, which has indeed been reported in some [27–29] but not all [30] histology studies based on human AD brain tissue. Another possible underlying cause of decreased CBV is capillary constriction. Recently, amyloid-β oligomers have indeed been shown to constrict capillaries in live human brain tissue, and localized capillary constriction was also observed in post-mortem fixed AD brain slices [31]. Changes in CBF have been associated with AD as well. A wealth of studies have shown this, where in the prodromal phase, CBF was shown to be increased, and decreased when clinical symptoms are overt (for a review see [3]). Similar capillary affliction as described above may underlie this decrease in CBF in cognitively impaired AD patients, but arteriolar dysfunction may also play a role. Reasons for the latter could be arteriolar CAA and/or atherosclerosis [9]. In mouse models of amyloidosis, it is ambiguous how CBV and CBF evolve. CBF has been shown to decrease [32–38], not change [36,39,40] or transiently increase [41] when compared to age-matched WT mice. Studies with some models show reduced capillary density [42,43], while studies with other models show preserved capillary density [40,41,44]. Regarding the diameter of the microvasculature, both capillary constriction [31] and capillary dilation [41] have been reported. An important flipside of the wide variety of models available is thereby illustrated: results obtained in one mouse model are not generalizable to other models, let alone to patients. On the other hand, the different models may help us understand the possible consequences of different types of amyloid-β accumulation. Interestingly, all studies that report no change or transiently increased CBF are performed with a model with a PS1 insertion [36,39–41]. The question then arises whether the PS1 insertion by some means protects the cerebral vasculature. Important features of models with an additional PS1 insertion are a shift in the amyloid-β-40:amyloid-β-42 ratio towards the more hydrophobic amyloid-β-42, as well as early and aggressive parenchymal plaque development [45]. An increase in hydrophobicity of the amyloid-β monomers has been hypothesized to lead to increased hydrophobicity of the oligomers, resulting in increased binding to cell membranes and increased neurotoxicity [46]. In the same way, it could be hypothesized that the increased hydrophobicity of oligomers reduces the movement of oligomers from the parenchyma to the vasculature, thereby decreasing the constrictive effects of amyloid-β on the vasculature. Indeed, in [36], where three amyloidosis models were directly compared, two models with CAA and without PS1 insertion showed a CBF decrease, and one without CAA and with PS1 insertion did not show a decrease in CBF.

There is much less data available on the relation between ATT and AD. One study in patients shows ATT is preserved in AD patients [47], whereas another shows a regional ATT increase [48]. To the best of our knowledge, no pre-clinical studies exist with amyloidosis mouse models and ATT. But given the lack of differences in mCBV and CBF, it is not surprising that no differences were found in ATT.

From the brain regions analyzed, the mCBV values found were highest in the thalamus and lowest in the white matter, which is in agreement with other studies [49–51]. There was however a large inter-animal variation in absolute mCBV values. For example, the mCBV values in the thalamus were between 3 and 6 % in WT mice, which is unlikely due to biological variation only. Some of this high variation may be due to varying quantities of lectin that effectively arrived in the brains of the different mice after the 200 μL i.v. injection, thereby creating differences in vessel wall labeling efficiency between animals. Correcting for such differences is complicated, as it is difficult to disentangle vessel wall labeling efficiency differences from true blood volume differences. However, since amyloid-β pathology is present in certain brain regions only, blood volumes could be normalized to un-affected brain regions such as the hypothalamus or cerebellum. This did not change the outcome however, as the APP/PS1 and WT mice still showed similar mCBV values after normalization to the hypothalamus or cerebellum (data not shown).

Despite the large volumes of tissue that were imaged, only a small number of biological replicates was used, which is the largest limitation of this study. The absence of significant differences between APP/PS1 mice and WT control mice suggests that parenchymal amyloid-β plaques have little or no effect on the structure and function of the cerebral vasculature, at least in mice. However, trends could be observed towards lower mCBV and cortical CBF in the APP/PS1 mice. If these were true, and not due to noise and/or natural variation, they could become significant with larger group sizes.

The size of the trend towards lower cortical CBF in this study (10-20 %) is however not on par with the observed CBF decrease in the clinic (42 %)[52]. Additionally, one important difference between this mouse model and AD patients is the lack of cerebral atrophy, despite the high amyloid burden. As atrophy is often hypothesized to contribute to cerebral hypoperfusion [53], our observations do not discount this possibility.

Furthermore, the age- and gender distribution was not fully equal between the groups, where the WT group contained 1 more male and was 1 month older in the microscopy study, and 2 more males and 2 months older in the ASL-MRI study. Preferably, age and gender would have been completely equal. Indeed, gender is known to influence cerebrovascular factors such as stroke risk and outcome [54,55]. Furthermore, female mice have shown higher rates of amyloid-β accumulation [56]. However, baseline brain perfusion and blood volume have been reported to be similar in male and female mice [55,57], and all transgenic mice showed advanced amyloid-β pathology, making it unlikely that our data was heavily biased by the gender difference. Also, brain perfusion has been shown to be stable in wild type adult mice [22,36], reducing the likelihood of an age bias. Moreover, there was no trend observable in the data towards lower or higher mCBV, CBF or ATT with age or gender (not shown). Another limitation of this study is the relatively low resolution used, which might not be sufficient to detect small constrictions in capillary diameter, especially if these were to be focal constrictions. It should also be taken into account that red blood cells could be hindered by such focal constrictions, lowering the local hematocrit, but leaving the mCBV, CBF and ATT unchanged. Further research could include imaging small areas at higher resolution to be able to detect such small changes. Also, despite the 3 to 4 mm thickness of imaged tissue, only 0.5 mm was used for mCBV quantification, due to dropping SNR values with deeper tissue areas. Possibly, further optimization of the tissue clearing and/or imaging could improve the SNR in deeper tissue and thereby improve the mCBV mapping. In our hands however, changing the clearing method to a method that results in tissue with a lower refractive index (RI) (*e*.*g*. ScaleA [58] or PACT [59]) did not give better results, due to extensive tissue expansion (data not shown). The expansion strongly drives up image acquisition times, whereas effectively, similar sizes of tissue are imaged when corrected for the expansion. Changing to a clearing method resulting in higher tissue RI such as 3DISCO [60] would be sub-optimal given the RI of the microscope objective used here and would require the use of solvents that are not compatible with the objective.

In summary, tissue clearing followed by microscopy of the brain vasculature and *in vivo* ASL-MRI was used in this study to measure cerebrovascular morphology and function in the APP/PS1 mouse model of amyloidosis with high parenchymal plaque burden. No changes were measured with either of the two imaging modalities, indicating that structure and function of the in the mouse brain vasculature are not heavily affected by the presence of amyloid-β plaques. The observed cerebrovascular dysfunction in AD patients may therefore not be much related to the presence of amyloid-β plaques, but originate more from other amyloid-β species such as oligomers and CAA, and/or from other non-amyloid-β-related factors.

## ACKNOWLEDGEMENTS

This research was funded by NWO, the Dutch National Science Organization, with the Innovational Research Incentives Scheme Vidi (L. van der Weerd) and by the heart-brain axis consortium of Cardiovasculair Onderzoek Nederland Society (CVON), part of the Dutch heart foundation (‘Hartstichting’).

