## Supplementary files for "Multi-scale Assessment of Brain Blood Volume and Perfusion in the APP/PS1 Mouse Model of Amyloidosis"

**Supplementary figure 1:** Oblique optical sectioning of the microvascular Cerebral Blood Volume (mCBV) maps, showing an axial gradient in mCBV values.

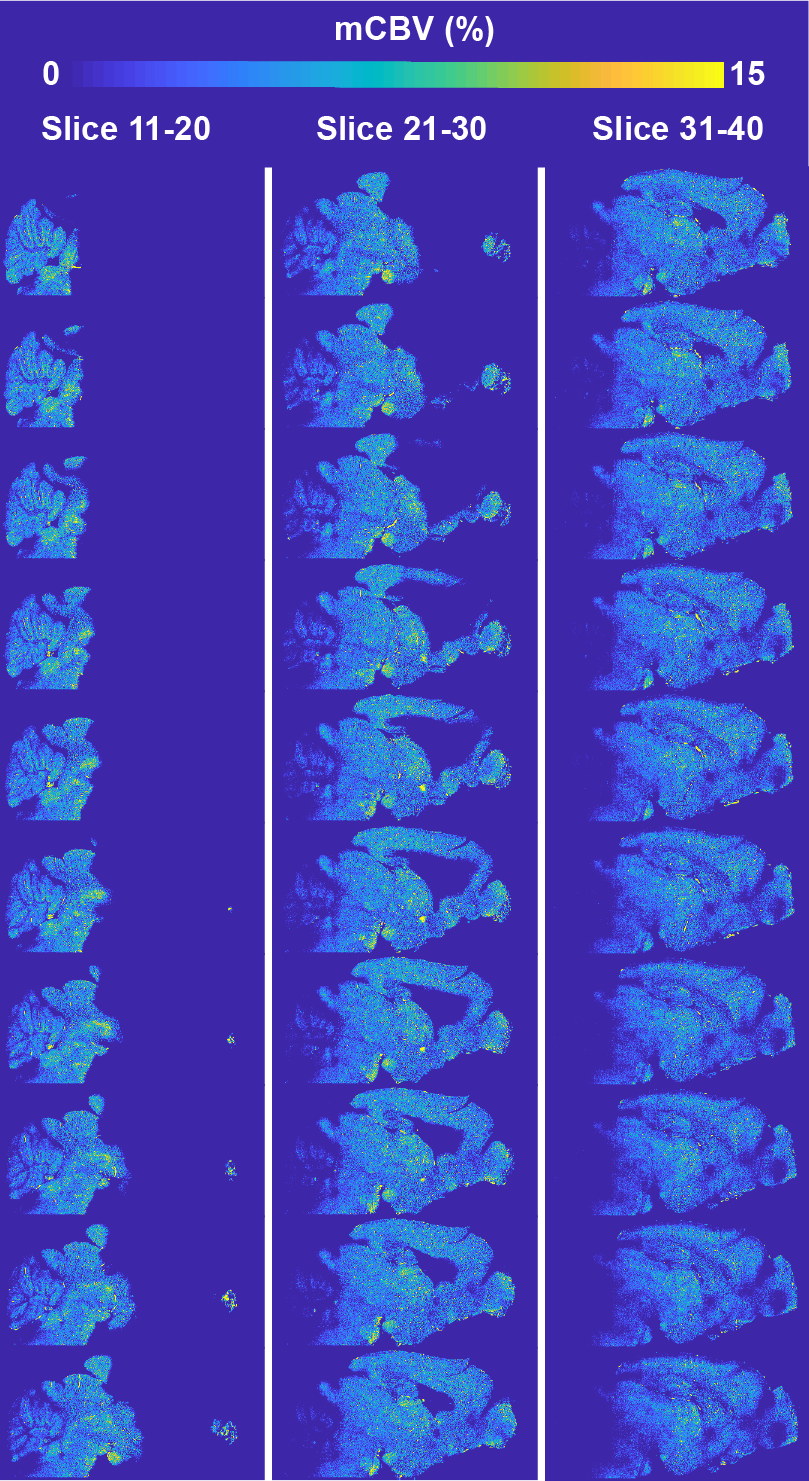

**Supplementary figure 2:** Aligned microvascular Cerebral Blood Volume (mCBV) maps, showing decreasing mCBV values with increasing imaging depth.

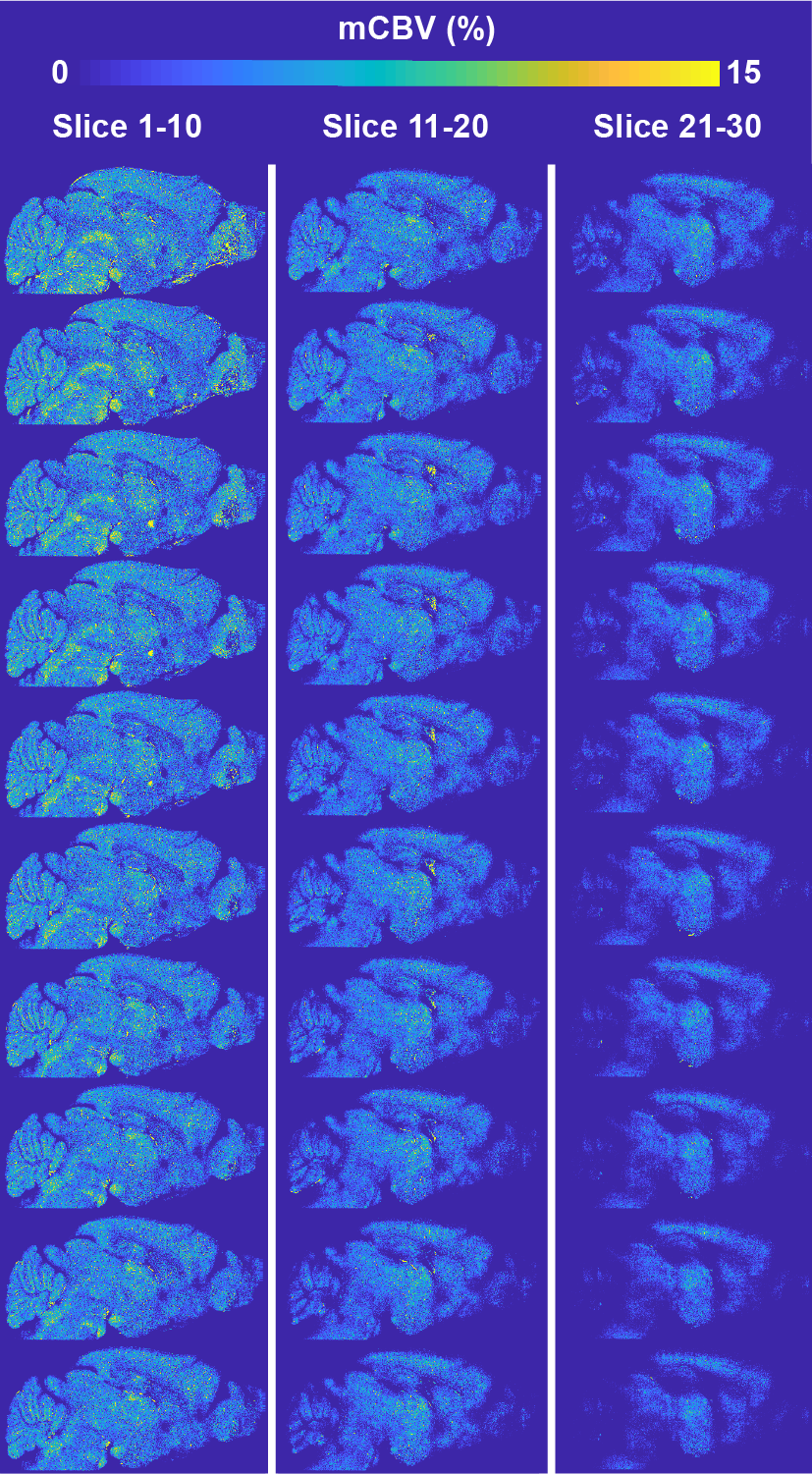

**Supplementary figure 3:** Examples of preserved vascular architecture around plaques. The yellow arrowheads point to vessels that are in the close vicinity to plaques, but seem unaffected by them.

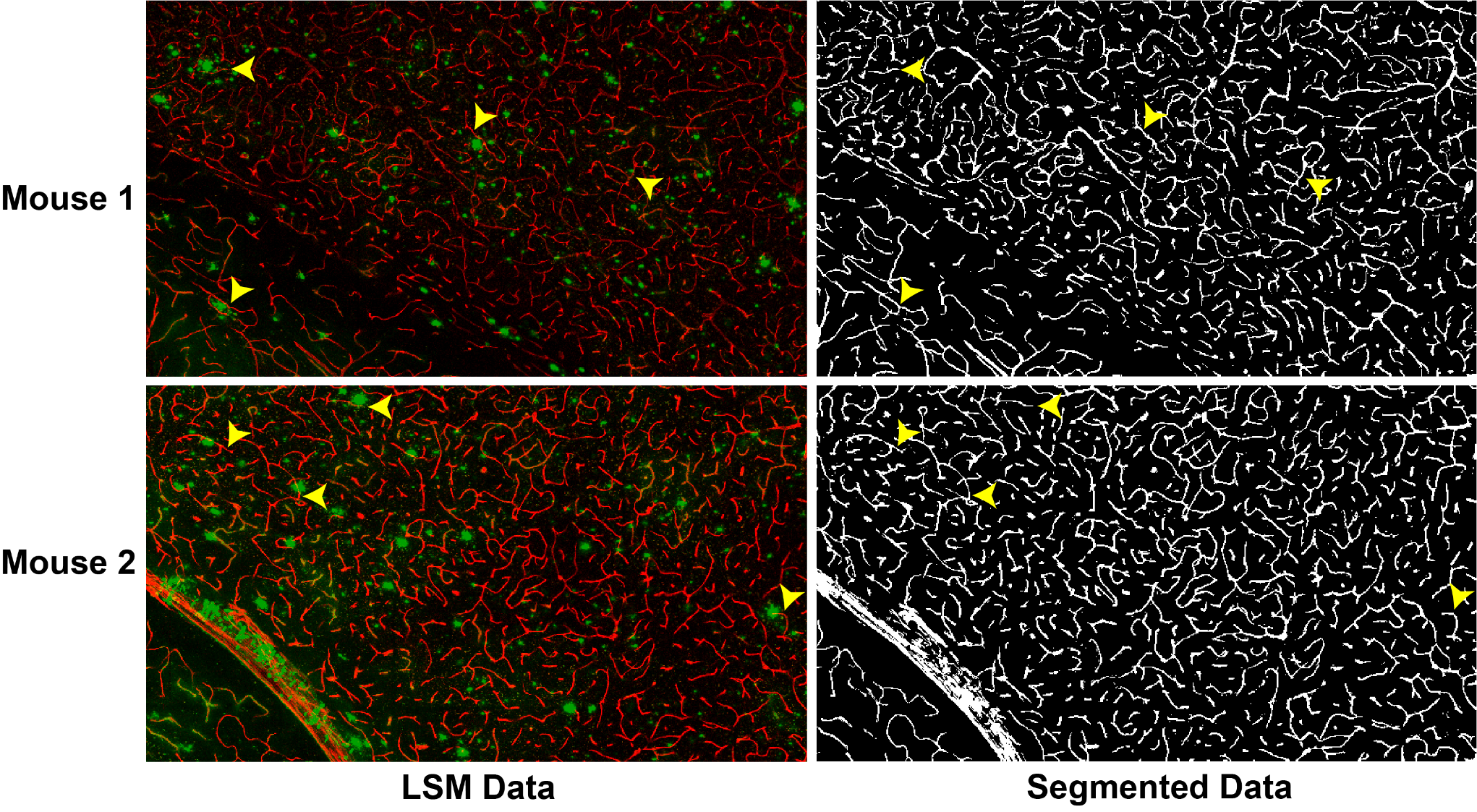
